# Ecological and evolutionary consequences of viral plasticity

**DOI:** 10.1101/289116

**Authors:** Melinda Choua, Juan A. Bonachela

## Abstract

Viruses can infect any organism. Because viruses use the host machinery to replicate, their performance depends on the host physiological state. For bacteriophages, this host-viral performance link has been characterized empirically and with intracellular theories. Such theories are too detailed to be included in models that study host-phage interactions in the long term, which hinders our understanding of systems that range from pathogens infecting gut bacteria to marine phage shaping present and future oceans. Here, we combined data and models to study the short- and long-term consequences that host physiology has on bacteriophage performance. We compiled data showing the dependence of lytic-phage traits on host growth rate (viral phenotypic “plasticity”) to deduce simple expressions representing such plasticity. We included these expressions in a standard host-phage model, to understand how viral plasticity can break the expected evolutionary trade-off between infection time and viral offspring number. Furthermore, viral plasticity influences dramatically dynamic scenarios (e.g. sudden nutrient pulses or host starvation). We show that the effect of plasticity on offspring number, not generation time, drives the phage ecological and evolutionary dynamics. Standard models do not account for this plasticity, which handicaps their predictability in realistic environments. Our results highlight the importance of viral plasticity to unravel host-phage interactions, and the need of laboratory and field experiments to characterize viral plastic responses across systems.

## Introduction

In the last three decades, advances in virology have unveiled the key role of viruses in a multitude of different ecosystems. From microbiota to the open ocean, viruses can affect almost any trophic level, showing a remarkable variety of life strategies that are shaped by viral reproductive mode and life-history traits. Let us focus on viruses that infect bacteria (bacteriophages). The cell lysis induced by marine phages, for example, removes up to 40% marine bacteria, thus enabling the remineralization of the resulting particulate and dissolved organic matter and, therefore, introducing alternative nutrient paths in the microbial loop (Abedon et al., 2009; Breitbart, 2012; Fuhrman, 1999). Phages can also alter the bacterial part of the human microbiota, as they can infect biofilms (Abedon, 2011) including gastrointestinal bacteria. Lytic phages have been suggested as an alternative to antibiotics (phage therapy) (Weld et al., 2004), and more generally to treat bacterial infections in plants and animals of commercial interest for the food industry (see (Golec et al., 2014) and references therein). Thus, understanding the dynamic interaction between host and phage in the short and long term is essential to make reliable predictions of high biological, medical, and commercial importance.

For obligately-lytic phages, the infection cycle starts when the virus encounters and attaches to the host cell (Calendar and Abedon, 2006). The phage then perforates the cell’s membrane and injects its genome into the cytoplasm. During early transcription, the first produced viral components divert the host synthesis machinery, inhibiting host replication. Host DNA is degraded and host components (nucleotides, ribosomes, ATP, etc) are used for the replication of the viral genome and synthesis of proteins that will compose the viral offspring. Late proteins, mostly structural, are assembled to compose the new virions, which are released when the so-called holin gene is expressed to facilitate host membrane lysis. This lytic cycle defines the main viral traits (Weinbauer, 2004): *i*) adsorption rate, or rate of successful encounters between host and virus; *ii*) eclipse period, or time between adsorption and the assembly of the first virion; *iii*) maturation rate, or rate of virion assembly; *iv*) latent period, or time between adsorption and cell lysis; and *v*) burst size, or offspring number. The adsorption process and rate depend on multiple factors, from the turbulence of the medium (Berg and Purcell, 1977) to phage morphology and host receptors (Schwartz, 1976). Although the degree of dependence of the eclipse period on the host differs across phages (e.g. T4 uses host RNA polymerase whereas T7 also uses its own), in all cases host ribosomes are used for viral protein synthesis (Calendar and Abedon, 2006; Walsh and Mohr, 2011). Virion assembly and DNA packaging rely on host ATP as energy source (Calendar and Abedon, 2006), host ribosome number and elongation rate (You et al., 2002), and host metabolic rates (Hadas et al., 1997). On the other hand, although it is not clear what determines the timing of lysis, the timing of the holin gene expression initiating this process is known to depend on both phage and host (Abedon et al., 2001). This timing influences not only viral generation time but also the final burst size (Gnezda-Meijer et al., 2006), relationship traditionally formulated as a trade-off by which larger offspring numbers require longer latent periods (Wang, 2006). Deeply affected by the maturation rate, this number of virions per infection can show a wide range of values and therefore is not the result of mere host nucleotide recycling (Brown et al., 2006; Maat et al., 2016; You et al., 2002).

Phage performance depends, in consequence, on host performance, e.g. host population number and physiological state (Wang et al., 1996). The latter has been documented with experiments that follow one single infection cycle (one-step growth), mainly for latent period or burst size, with a variety of hosts in a diversity of environments ((Abedon, 1989; Abedon et al., 2003; Gnezda-Meijer et al., 2006; Golec et al., 2014; Kokjohn et al., 1991; Maat et al., 2016; Middelboe, 2000; Proctor et al., 1993; Webb et al., 1982), to name a few). In all cases, an improvement in the host physiological state led to a shortening of the latent period but an increase in burst size. Comprehensive studies that explore simultaneously how all the viral traits described above are affected by the host physiological state are, however, difficult to find (Hadas et al., 1997; You et al., 2002).

The experiments above focus on the growth rate at the moment of infection to represent host “quality” (Wang et al., 1996), and show that viral trait values change in the short term with changes in the host, effectively the most important component of the virus’ environment. Hereby, we will refer to such changes as viral phenotypic plasticity. This plasticity is, however, typically ignored in phage studies. For example, most experiments compiling information about phage traits are conducted under ideal conditions for the host (that is, maximal rates for host growth, see discussion in, e.g. (Hadas et al., 1997)). In nature, however, the situations in which the host grows at its maximum rate are more the exception than the rule. As a consequence, theory or field conclusions built on such trait values are potentially biased, including viral count and effect on the microbial community structure, population density, and dynamics.

From a theoretical point of view, intracellular descriptions replicate single-infection-cycle data (You et al., 2002), but their level of detail renders these models computationally expensive and difficult to parametrize (Birch et al., 2012), and therefore impractical for the study of the long-term behavior of any specific host-virus system. Ecosystem models, for instance, rarely include viruses or they are included (due to computational constraints) with simplified terms that neglect plasticity (Mateus, 2017). On the other hand, optimal (i.e. fitness maximizing) latent periods have been estimated using fixed viral traits and/or fixed host concentrations accounting for host quality (Abedon et al., 2001; Wang et al., 1996), although decoupling host growth rate and density precludes these calculations from predicting the dynamics of the system. Dynamic-model attempts to include plasticity used different fixed trait values for different host growth rates (Middelboe, 2000), or case-specific effective expressions (Rabinovitch et al., 2002) aimed to improve the design of one-step growth experiments (Aviram et al., 2015).

Therefore, past experiments and theories advanced knowledge on how the host growth rate affects phage development; however, how such a link affects the ecological and evolutionary dynamics of the phage-host system remains largely unknown. Here, we aim to fill this knowledge gap by addressing the following questions: how does viral plasticity affect the eclipse period, maturation rate, latent period, and burst size in the short and the long term? How do these trait changes affect the ecological interaction between host and phage in static and dynamic environments?

To answer these questions, we focus here on T phage infecting *Escherichia coli*, one of the most common host-phage model systems. We first compiled the available data on T-phage trait changes motivated by host growth, to deduce and assess the generality of our own functional forms linking host growth rate with eclipse period and maturation rate. Because the factors that trigger lysis are unknown, we assumed that latent period and associated burst size are evolutionary outcomes. We then studied such emerging evolutionary strategies by embedding the data-deduced expressions in a standard host-phage mathematical model, which allowed us to study the short- and long-term ecological and evolutionary behavior of the system. Our results provide insight on the mechanisms underlying the timing of lysis under a diversity of environmental conditions, including steady states and dynamic nutrient changes, and how plasticity affects such timing and burst size. Our expressions can help improve the predictability of host-phage models, from small-scale to earth-system models. Moreover, our findings have the potential to be generalized to any lytic phage, including viruses that affect biofilms or used for phage therapy or industrial setups, and motivate novel experiments aimed at characterizing viral plasticity across systems.

## Methods

### Data

We compiled data from one-step growth experiments (Ellis and Delbrück, 1939) that focused on measuring the viral trait values for various host growth rates (Birch et al., 2012; Golec et al., 2014; Hadas et al., 1997; You et al., 2002) (see table 1). Specifically, these experiments followed one infection cycle of *E. coli* by either the T7 (Birch et al., 2012; You et al., 2002) or T4 phage (Golec et al., 2014; Hadas et al., 1997), both obligately lytic. All the experiments used chemostats in which host growth rates were regulated by either using different sources of carbon (Birch et al., 2012; Hadas et al., 1997) or using different dilution rates (Golec et al., 2014; You et al., 2002); culture and infection temperatures were kept to 30*C* in (You et al., 2002), whereas the rest used 37*C* (see detailed description in the online Appendix, section A). In all cases, the eclipse period, *E*, and maturation rate, *M*, were either reported or the original experimental data provided. The latter allowed us to estimate *E* and *M* using their standard definition (time after infection at which the first virion is assembled, and rate of increase of the chloroform-generated PFU data, respectively).

**Table 1:** Compilation of data from the literature.

| Source | Host | Phage | Temperature | Data for fits |
| --- | --- | --- | --- | --- |
| (You et al., 2002) | *E. coli* | T7 | 30C | Extracted from original data |
| (Birch et al., 2012) | *E. coli* | T7 | 37C | Extracted from original data |
| (Hadas et al., 1997) | *E. coli* | T4 | 37C | From reported *E* and *M* data |
| (Golec et al., 2014) | *E. coli* | T4 | 37C | Extracted from original data |

The first and last databases used different dilution rates to vary the host growth rate, whereas the central two databases used different sources of carbon.

#### Data analysis

We fitted the extracted data for the eclipse period and maturation rate as a function of the host growth rate, *µ*, to obtain functions that include plasticity, *E*(*µ*) and *M* (*µ*). To facilitate comparison across experiments, we normalized *µ* using the maximum growth rate (*µ*_*max*_) reported in each experiment for host optimal conditions. Such values were compatible with tabulated values for *E. coli* maximum growth rates at the temperature used in each case (Herendeen et al., 1979).

For the fits, we focused on simple functional forms that can replicate the data across datasets and are also biologically meaningful. Simple decreasing exponentials, for example, match qualitatively with a reduced number of parameters a negative correlation with host growth rate reaching a lower plateau for high *µ*. As shown in Figs.1, A1–A3 (left panels) and table A1, the data available from all our sources are explained separately by:

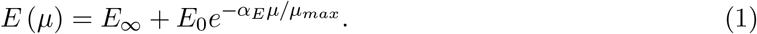

**Figure 1:**
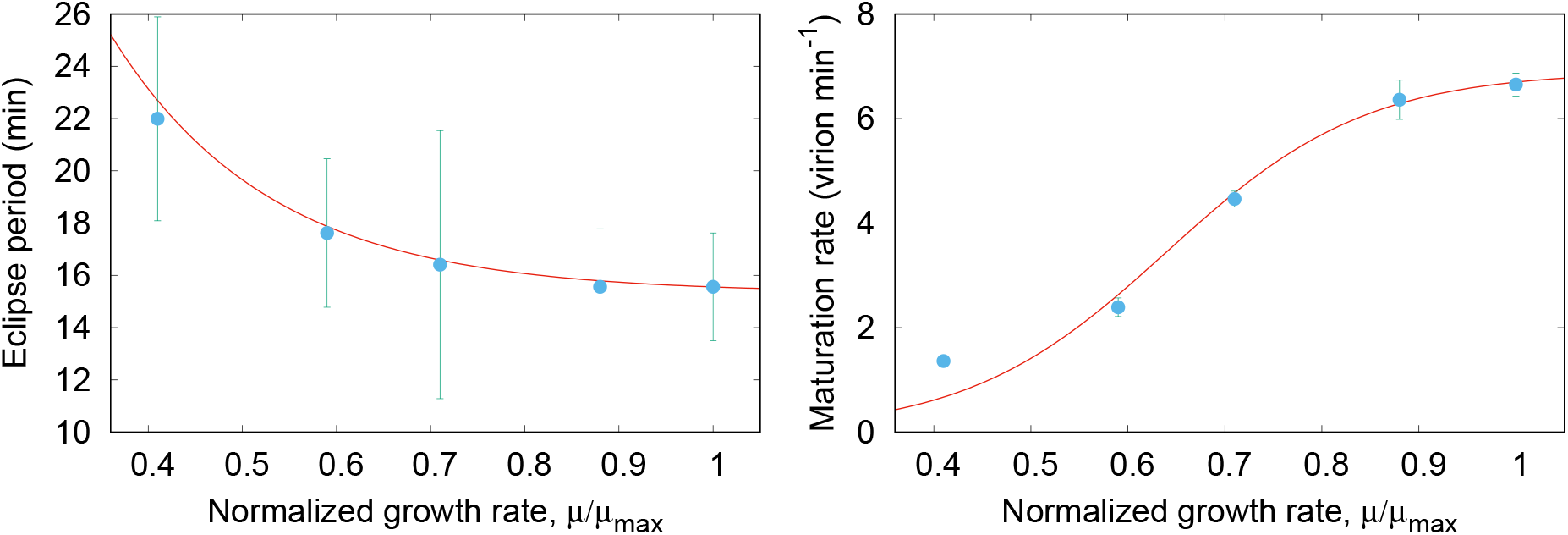
Data extracted from the original one-step growth experiment data in (You et al., 2002), and proposed curves. Left: Eclipse period, *E*; Right: Maturation rate, *M*.

This function captures the high *E* values observed for very low growth rates, with *α*_*E*_ determining how steeply the function decreases towards the minimum-value plateau, given by *E*_∞_ and *E*_0_. Table A1 shows the parameter values across examples, as well as indicators of the closeness of the function to the data (adjusted *R*^2^ and root-mean-squared deviation). On the other hand, the maturation rate data show an early slow growth that accelerates for intermediate growth rates and ultimately saturates (Figs. 1, A1–A3, right panels), which resembles a sigmoid function such as:

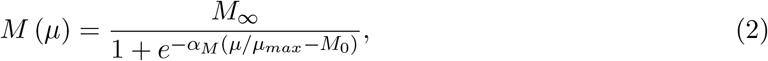

where *M*_∞_ represents the upper plateau, *α*_*M*_ how steeply *M* grows to that plateau, and *M*_0_ the location of the function.

Eq.(1) and Eq.(2), thus, represent well the plastic response observed with different T phages and temperatures, sources of variation that are encoded in the various coefficients (tables 1 and A1). When parametrizing the model below, we will focus on the (You et al., 2002) database (i.e. T7 phage infecting *E. coli* at 30*C*), the most complete dataset, and the *E* and *M* expressions deduced for this case. Although other qualitatively-similar functions may also fit the data (see section B), our results are robust against the specific functional forms for *E*(*µ*) and *M* (*µ*) (see below).

### Theoretical model

To represent the dynamics of the host-phage system, we used a classic model that includes explicitly the delay between infection and lysis (Levin et al., 1977). This model has proven to be realistic from both the ecological and evolutionary points of view (Bonachela and Levin, 2014), and keeps track of the dynamics of free host cells, [*C*], infected hosts, [*I*], and free viruses, [*V*], with the following delayed differential equations:

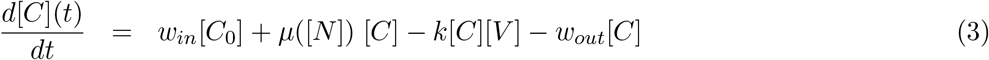

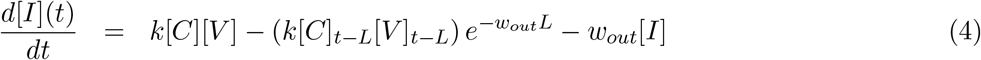

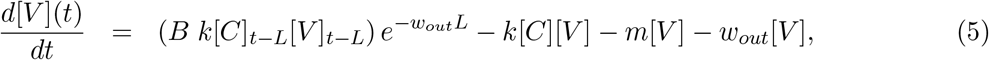

where *B* represents the burst size and *L* the latent period (see symbols and units in table A2). We emulate here a two-stage chemostat, a setup in which a first chemostat allows for a virus-free culture of the host to reach a controlled stationary density, [*C*_0_], which feeds the second chemostat where the interaction with the phage, Eq.(3)–(5), happens (Husimi et al., 1982). Such a setup reduces significantly the region of the parameter space for which the typical (predatory-prey-like) oscillations are expected, and reduces evolutionary pressures on the host thus allowing us to focus on the evolution of the phage only. The inflow of fresh hosts at a rate *w*_*in*_ is represented in the first term of Eq.(3); the second term represents the growth of the host, whereas the third term represents the new viral attachment and infections; the last term represents the host removal at a rate *w*_*out*_ during dilution events in the chemostat. New infections increase [*I*] (first term in Eq.(4)), which can disappear during dilution (last term) or due to the lysis of cells that became infected exactly a latent period in the past (second term, where 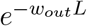 is the survival probability for infected cells during the extent of the latent period). The virions released by such cells at lysis contribute to the pool of free phages (first term in Eq.(5)) which, in turn, can disappear after infecting free hosts (second term) or due to dilution or natural mortality (last two terms). We assume that the growth rate for free hosts can be described by the simple Monod model (Monod, 1950), *µ* = *µ*_*max*_[*N*]/([*N*] + *K*_*N*_), where *K*_*N*_ is the half-saturation constant for growth (inversely correlated with growth affinity), and [*N*] is the concentration of the nutrient in the second chemostat, which changes with time according to the equation:

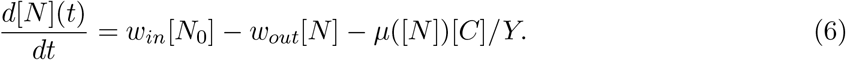

[*N*_0_] is the input concentration of nutrient entering this second chemostat, and *Y* the yield parameter that accounts for the host efficiency to transform uptake into growth. See sections C and D for further details and assumptions.

The equations above define the ecological dynamics. Because the adsorption rate, *k*, can be affected by multiple factors not considered here (e.g. host plasticity and evolution), we assume a constant *k* for all phage phenotypes in our one-host system. From the remaining viral traits, *E* and *M* are univocally determined by the host growth rate; *B* is set by the maturation rate and the time between the end of the eclipse period and lysis, relationship typically represented by the linear function *B* = *M* [*L* − *E*] (Wang, 2006). This expression assumes that only lysis and not intracellular resources limit the offspring number. Considering viral plasticity, the burst size can be rewritten as:

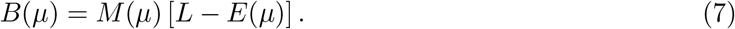

The latent period, only remaining trait, characterizes therefore the viral phenotype (indistinctly the holin gene, responsible for initiating lysis, characterizes genotypes). We will assume that *L* is the only adaptive trait, i.e. the end of the lytic cycle is an emergent, evolutionary outcome.

#### Analytical approach

To obtain an initial understanding about the effect of viral plasticity on the long-term behavior of the system above, we analyzed its ecological and evolutionary stationary states. For the ecological stationary states, we calculated the solutions to the equations *d*[*C*]/*dt* = *d*[*I*]/*dt* = *d*[*V*]/*dt* = *d*[*N*]/*dt* = 0 and studied the associated stability (see C-D and (Bonachela and Levin, 2014)). For the evolutionary stationary state, we conducted invasion analyses by studying the stability of a phage-host system that, after reaching equilibrium, is inoculated with an invading viral phenotype challenging the resident phage. Such invaders can also represent new genotypes that result from mutations in the holin gene and therefore show different latent periods. The invasion analysis determines the phenotype (i.e. *L*) that maximizes fitness, the Evolutionarily Stable Strategy (ESS) (Dercole and Rinaldi, 2008; Geritz et al., 1998). The ESS can be identified as the phenotype that will dominate the system in the long term even in the presence of all possible variability. See section E for further details.

#### Numerical approach

The analytical approach above requires important assumptions such as the quick relaxation to the ecological equilibrium before mutants enter the system, which risks overlooking the dynamic influence of plasticity. Under realistic conditions, plasticity and evolution can interact non-trivially, which may in turn feed back to the ecological dynamics of host and phage (Lennon and Martiny, 2008). Such dynamic interaction, however, precludes the analytical calculation of a closed expression for the ESS. To go beyond such constraints, we used a numerical eco-evolutionary simulation framework in which both ecology and evolution occur at the same timescale (Bonachela and Levin, 2014; Lomas et al., 2014; Menge and Weitz, 2009).

Starting from a single viral phenotype and host populations, Eqs.(3)–(7) are integrated thus providing the ecological dynamics of the system. Mutants originate from parental phenotypes, at random times, by means of a roulette-wheel genetic algorithm in which the most abundant phenotype shows the highest probability to mutate (Menge and Weitz, 2009). Thus, multiple resident and mutant phenotype populations compete for the single host, with mutation and natural selection facilitating an alternation of dominance until, eventually, a phenotype rises that cannot be challenged by any other mutant. Such a strategy (the ESS) can be identified as an evolutionary stationary state in the representation of the dominant *L* against time. See F for details.

With such a framework, we ran *in silico* experiments aimed at finding the long-term evolutionary behavior of our system. Motivated by the viral plasticity experiments compiled here, we varied the host growth rate: *i)* using different input concentrations, [*N*_0_]; *ii)* using different sources of sugar, which we emulated by using a variety of growth affinities (half-saturation constants, *K*_*N*_, see above); or *iii)* using different dilution rates, *w*_*out*_. We also explored the evolutionary outcome of scenarios away from stationarity. To this end, we repeated the experiments above implementing a rapid variation of the environment by allowing the input concentration to vary on a daily basis by a random factor up to *σ*, with *σ* = 0.3 and *σ* = 0.5.

In all cases, to quantify the trait ranges enabled by plasticity we normalized the resulting trait curves by their minimum value. Such normalization also facilitates comparison across the methods described above to control *µ*, and with the ESS predicted by a non-plastic version of the model (*L*_*non*_ and *B*_*non*_). Because non-plastic models would use fixed trait values typically obtained from available experiments, which standardly set optimal conditions for the host, we assumed for this comparison that *L*_*non*_ and *B*_*non*_ correspond to the *L* and *B* values obtained for the maximum *µ* observed in the focal simulation (see section E).

A last *in silico* experiment gauged how plasticity alters predictions on host-virus interactions in diverse dynamic scenarios with sudden changes in nutrient availability. In the two-stage chemostat represented by Eqs.(3)–(7), with a baseline nutrient input concentration that sets the host growth rate to approximately half of its maximum, we explored three consecutive situations: a sustained nutrient pulse that takes the host to its maximum growth rate and another that takes the host to 2/3 its maximum, followed by a reduction in the input concentration that decreases host growth to 1/3 its maximum. We fixed the duration of each event to 20 days. With this setup, we compared the predictions of three different versions of the model above: *i)* one that ignores phages (mimicking most of current ecosystem models); *ii)* a non-plastic description of the phage (like standard host-phage models, i.e. fixed trait values obtained by assuming optimal conditions for the host); and *iii)* our plastic description. For the latter, we used our analytical ESS expression for *L* (with *E*, *M*, and *B* changing dynamically), thus simulating the most dominant/representative phenotype of the focal region of the ecosystem.

Finally, we tried all the experiments above also with one-stage chemostats to test the role of the marked oscillations expected in such case.

## Results

### Analytical results

Two-stage chemostats facilitate stationarity in our system by reducing ecological oscillations and preventing a co-evolutionary arms race between host and virus. From the ecological point of view, the nontrivial stationary state in which both host and viral population coexist is given by Eqs.(A4)–(A6). As described in section E, the associated evolutionary stationary state (i.e. the ESS) is:

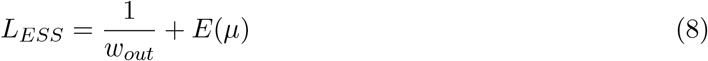

where *w*_*out*_ is contributed by the exponential term in Eqs.(4) and (5) and, therefore, represents more generally the removal rate for intra-cellular virions (equivalently, infected hosts). Eq.(8), together with the trade-off function Eq.(7), provides the associated burst size:

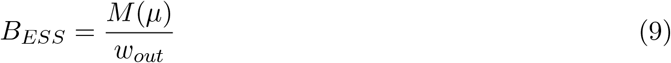

This ESS maximizes the viral fitness and minimizes the amount of hosts (i.e. resources) needed. Moreover, such a strategy is convergence-stable, as phenotypes closer to the ESS can always invade phenotypes further from *L*_*ESS*_ (section E). Note that the expressions for both the ecological and evolutionary stationary states are valid regardless of the specific details of *E*(*µ*) and *M* (*µ*).

### Numerical results

Due to the impossibility to find a closed form for the host growth rate as a function of time and the constraints of the analytical calculations, the diversity of scenarios below requires the use of the eco-evolutionary simulations. In order to have a reference curve from the analytical approach that we could compare with the numerics, we informed the analytical ESS with the numerical host growth rate. For all our simulations, we parametrized the model using table A2 and the parameters obtained from the (You et al., 2002) case. Note that, following standard practice for chemostats, we set *w*_*in*_ = *w*_*out*_ although, for coherence with the analytical calculations, we will keep referring to the dilution rate as *w*_*out*_.

#### Long-term behavior for different host growth rates

The ecological interactions including plasticity (Eq.(3)–(7) with Eq.(1)–(2)), together with evolution, give rise to trait dynamics such as those shown in Fig.A5 (right panel). Mutation and competition enable the exploration of the phenotypic space until, eventually, the ESS emerges and the system reaches an evolutionary stationary state. Because this exploration is intrinsically stochastic, the evolutionary stationary state values resulting from each replicate are spread around a mean value that we define as *L*_*ESS*_. Here, we used 300 replicates for each example.

We first studied the *L*_*ESS*_ emerging from such evolutionary dynamics using [*N*_0_] to control the host growth rate. Using input concentrations that ranged from [*N*_0_] = 7 × 10^−5^ *mol l*^−1^ to [*N*_0_] = 2 × 10^−3^ *mol l*^−1^, and fixing the dilution rate to *w*_*out*_ = 15 *d*^−1^, the stationary host growth rate ranged from ∼ 15.5 *d*^−1^ to ∼ 39.0 *d*^−1^, respectively (in Fig.2, expressed as fractions of *µ_max_*). Thus, there is a positive correlation between the host growth rate and nutrient input concentration. For reference, Fig.2 shows also the curve obtained using Eq.(8) (similarly for Eq.(9) for the the burst size). The maximum range of variation for *L*_*ESS*_ is approximately 8%, while there is a ∼ 18-fold range for *B*_*ESS*_. Latent period decreases and burst size increases with host growth rate, thus opposite to the usual positive correlation between infection time and offspring number (Fig.A8, left). This variation represents an improvement in phage performance with host growth rate, as increasing [*N*_0_] (i.e. *µ*) leads to an increased concentration of viral individuals and a consequent decrease in the concentration of hosts (Fig.A7, left). This relationship, together with the negative correlation between *L*_*ESS*_ and *µ*, results in an increase of the latent period as [*C*]_*st*_ increases (Fig.A7, right). The typical non-plastic description for the system shows the minimum possible *L*_*ESS*_ and maximum possible *B*_*ESS*_ within the range of variation of these traits (dashed line, Fig.2 and section E).

**Figure 2:**
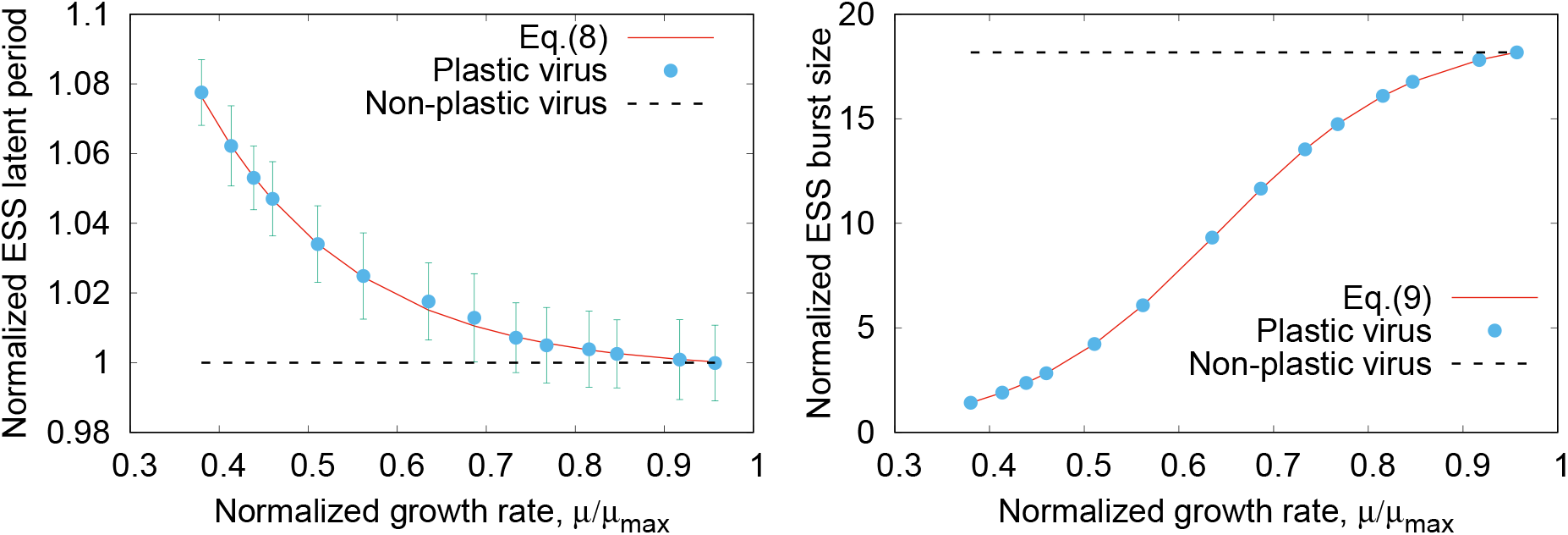
Evolutionary results from theory (solid line) and averaging over simulations (points) altering the host growth rate using [*N*_0_] = 7 *×* 10^−5^ *mol l*^−1^-[*N*_0_] = 2 *×* 10^−3^ *mol l*^−1^ with *w*_*out*_ = 15 *d*^−1^ and *K*_*N*_ = 9 *×* 10^−5^ *mol l*^−1^. The whiskers represent the standard deviation obtained across replicates, and the dashed line represents the value that typical models ignoring plasticity would use. Both vertical axes have been normalized using the minimum value obtained in the simulations. Left: Latent period, *L*. Right: Burst size, *B*.

We observed similar results when varying the host growth rate using different *K*_*N*_ values as a proxy for different sources of carbon (not shown). When using the dilution rate as a way to vary the host growth rate, however, varying *w*_*out*_ affects both terms from Eq.(8); moreover, in two-stage chemostats host growth rate and dilution rate are negatively correlated (illustrated theoretically in (Bonachela and Levin, 2014)). Consequently, both *L*_*ESS*_ and *B*_*ESS*_ are increasing functions of the host growth rate (Fig.3, and right panel in Fig.A8), with a very limited range of possible *µ*. Thus, this case shows a behavior similar to that of the non-plastic description, and respects the classic positive correlation between *L* and *B*, which sets an evolutionary trade-off between generation time and offspring (Fig.4, left, where both *L* and *B* show a very wide range of variation).

**Figure 3:**
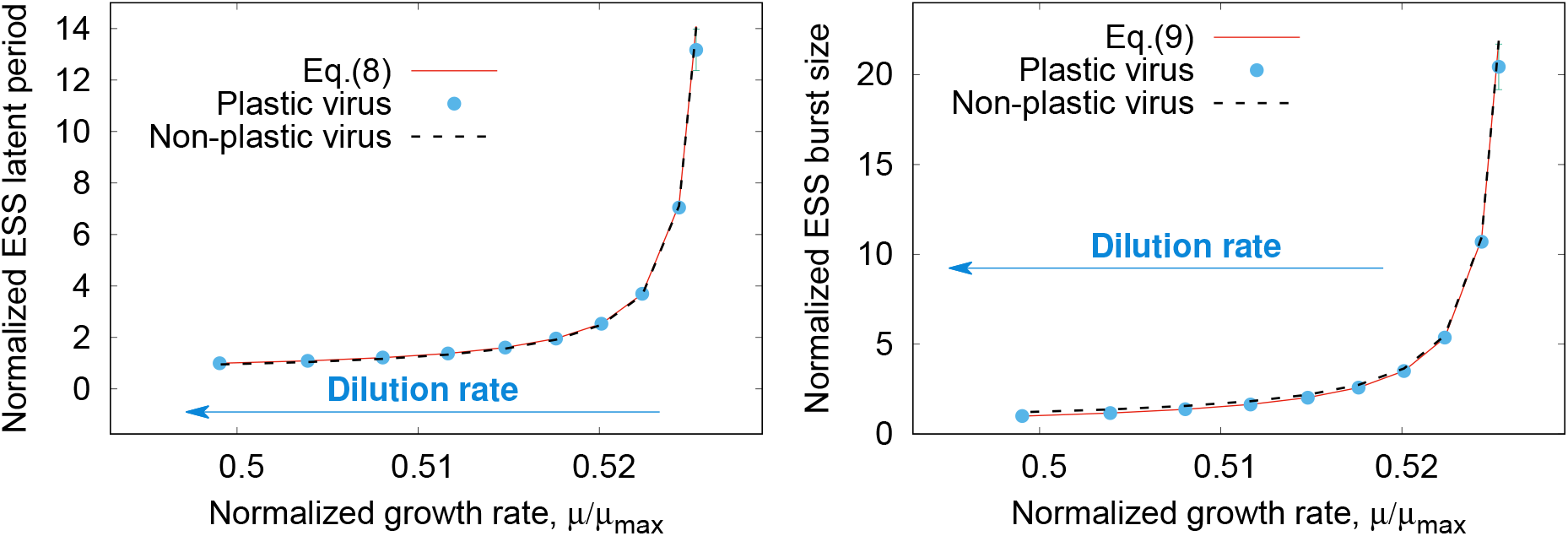
Evolutionary stable strategy obtained modifying the growth rate through *w*_*out*_ = 0.1*d*^−1^ to *w*_*out*_ = 0.91*d*^−1^ with [*N*_0_] = 1 *×* 10^−4^ *mol l*^−1^ and *K*_*N*_ = 9 *×* 10^−5^ *mol l*^−1^. Left: Latent period. Right: Burst size.

**Figure 4:**
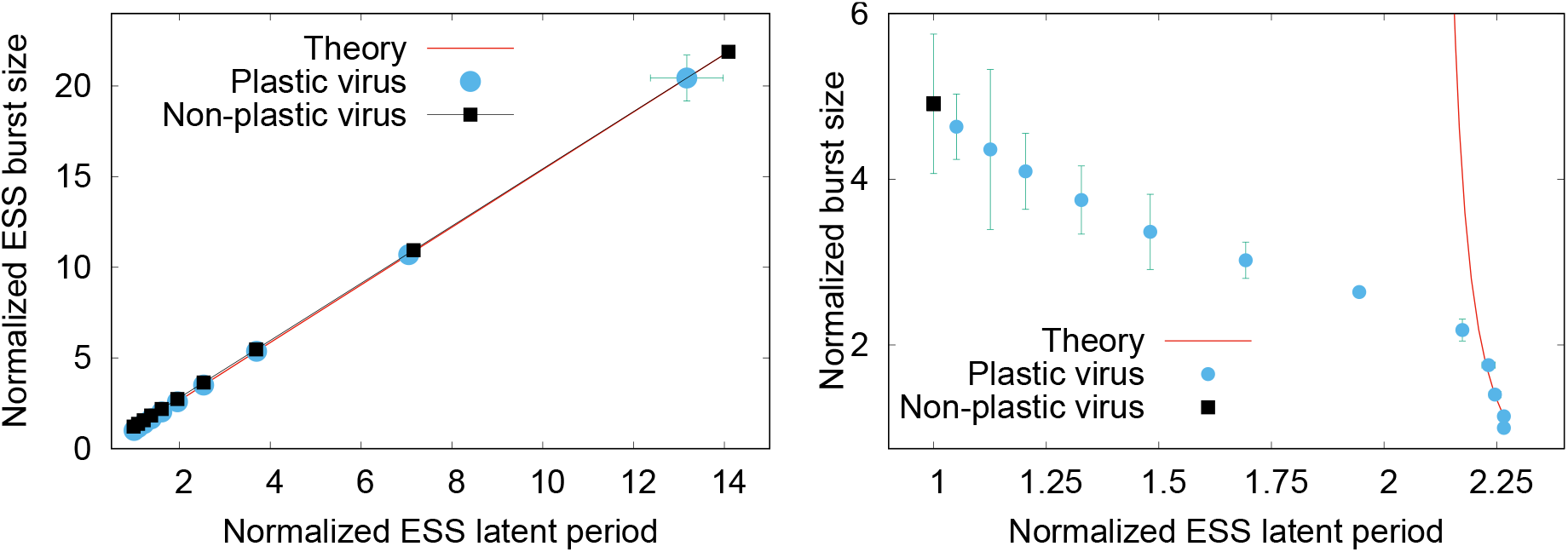
Left: Trade-off curve obtained when host growth rate is varied using different dilution rates. Right: Same curve obtained using different nutrient input concentrations, in this case using one single chemostat for the interaction phage-host (i.e. limit [*C*_0_] = [*N*_1*st*_] = 0 in our model), with *w*_*out*_ = 8.16 *d*^−1^.

#### The role of fluctuations

We repeated the three methods above with classic one-stage chemostats (i.e. assuming no inflow from the first chemostat). We observed qualitatively similar trends as above: a *L*_*ESS*_ that decreases and a *B*_*ESS*_ that increases with host growth rate (Fig.A6 left, and Fig.4 right), with the exception of the positive correlation between burst size and latent period obtained when using *w*_*out*_ to control *µ*. When an equilibrium was reached (in Fig.4 and A6, range *µ/µ*_*max*_ ∼ 0.25 − 0.35), the stationary state was accurately described by Eqs.(A4)–(A6) and, in consequence, the ESS was well described by our analytical expressions (solid line, Eqs.(8)–(9) informed by the stationary growth rate). These expressions, however, failed to describe the wide and oscillations found for the rest of input concentrations, for which a steeper decrease for *L* as a function of *µ* and a milder increase for *B* was observed (*µ/µ*_*max*_ > 0.35). Thus, the range of *L* increased with respect to the theoretical prediction, and that of *B*, decreased. On the other hand, fluctuations introduced in two-stage chemostats by means of a random (daily) forcing of the input concentration did not lead to significant departures from the analytical expression (Fig.A6, right).

#### Short-term behavior under dynamic environmental conditions

Finally, we shifted the focus to short-term population dynamics and compared the predictions of three different versions of the model for various dynamic events. As Fig.5 shows, the model without viruses shows a host population that follows the qualitative behavior of the nutrient input (see also Fig.A9). For the non-plastic description, the phage population shows the qualitative profile of the nutrient input as a response to the attempts of the (free) host population to cope with nutrient changes. However, the viral ecological pressure maintains the average host population around approximately the baseline level throughout the whole experiment. Different (fixed) phage parametrizations only altered the result quantitatively and, in some cases, introduced an oscillatory behavior for the different nutrient regimes (results not shown). Even in the presence of oscillations, nonetheless, the averages for each dynamic event of the phage and host populations consistently agreed with the behavior described above. With our plastic description, unexpectedly, events that increase the host growth rate lead to a decrease in the host population with respect to the baseline, linked to an increase in phage population (Fig.5, and Fig.A9 right). Also against intuition, nutrient scarcity leads to an increase in the host population and a decrease of the phage population. The qualitative behavior of the host population in the plastic description is, therefore, radically different from that of the virus-free and non-plastic versions. The behavior of the phage population is qualitatively similar to that shown by the non-plastic description agreeing quantitatively, in the case shown in Fig.A9, when the host growth rate reaches *µ* = *µ*_*max*_. On the other hand, it is possible to find population oscillations also with the plastic description, although phage acclimation (via plasticity) translates generically into damped oscillations when transitioning between regimes. We repeated the same experiment considering remineralization, which did not change qualitatively the described behavior (section G).

**Figure 5:**
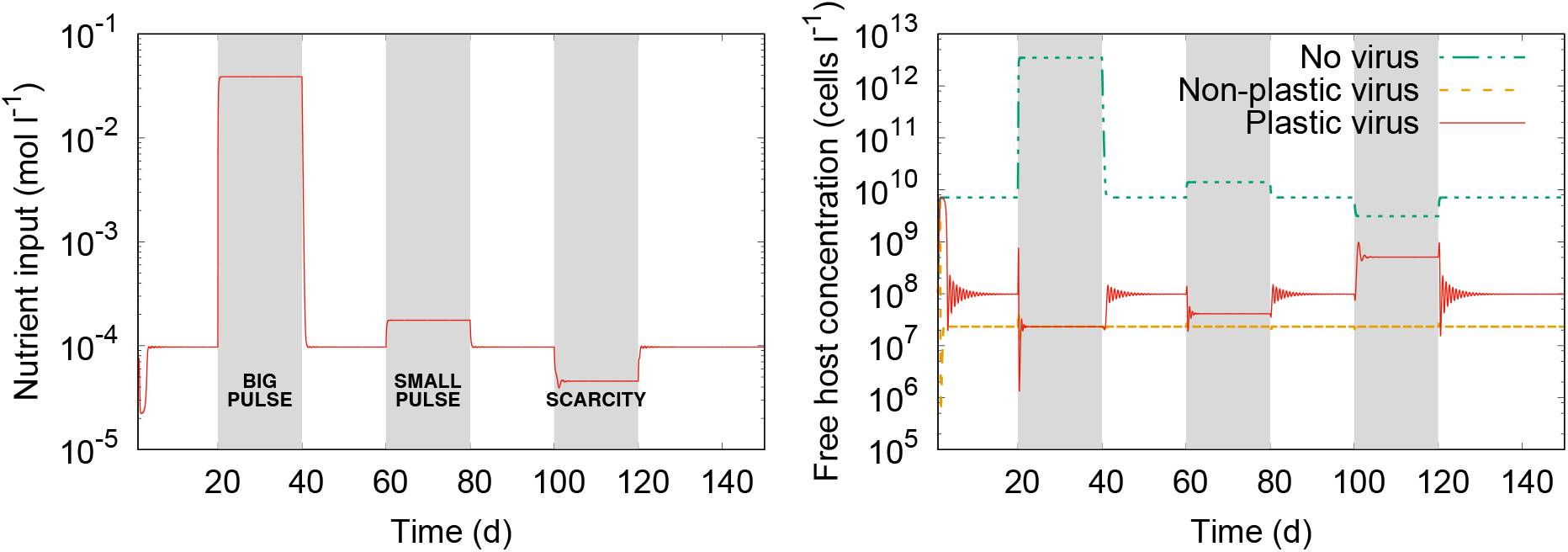
Dynamics of one replicate of the two-chemostat setup for three different events (shaded areas) in which the nutrient inflow brings the host growth rate to *µ* = *µ*_*max*_, *µ* = 2*µ*_*max*_/3, and *µ* = *µ*_*max*_/3. Left: Nutrient profile. Right: Total number of non-infected hosts.

## Discussion

We have used available data on how T-phage eclipse period and maturation rate change with host growth rate to study analytically and computationally the effects that such viral phenotypic plasticity has upon phage traits, and phage and host populations. To this end, we have assumed the latent period to be an emergent evolutionary outcome that, together with the eclipse period and maturation rate, determines the burst size.

### Generic mechanisms and functional forms for phage plasticity

The functional forms for *E* and *M* we introduced here consistently explain the data obtained for different T phages under different experimental conditions. The eclipse period data are compatible with a decreasing exponential function of the host growth rate, with a non-negligible minimum plateau. In addition, the maturation rate data show an increase similar to a sigmoid from a negligible value to reach a certain saturation value. Both qualitative forms are biologically meaningful. An increase in host growth rate correlates with an increase in the transcription and translation machinery and rates (as would happen in a healthy host), thus leading *E*(*µ*) to show the initial decline in the time needed to synthesize viral components. The lower plateau may relate to physiological limits to such synthesis, for example limits to host metabolic rates (Birch et al., 2012), or translation limitation due to emerging bottlenecks (You et al., 2002). For similar reasons, increasing host growth rates materialize in a quick improvement for *M* (*µ*), slowing down and eventually being limited at high growth rates by ribosome efficiency, polyribosome elongation rate, and late protein expression (Calendar and Abedon, 2006; You et al., 2002). This biological interpretation of the proposed functions is supported by the changes in phage strain and temperature across databases (section B).

Note that the mechanisms mentioned above as explanatory of the qualitative shape of our expressions are a fundamental part of the replication of most lytic phages, which suggests that qualitatively-similar curves may describe generically phage plasticity. Phage-specific details and influential environmental factors such as temperature are encoded in the parameters of such functions (table A1). On the other hand, because all the compiled data share the same host species, we cannot discern whether the host influences any parameter other than the normalization factor (i.e. *µ*_*max*_), which we used naively here to deal with temperature-related and host strain-specific differences. Nonetheless, our analytical results for *L* and *B*, which agree with the numerical results for all the non-oscillatory cases, do not depend on the shape of *E*(*µ*) and *M* (*µ*) functions. Exhaustive experiments aiming at gathering information on different phages and hosts are needed in order to assess the generality of the functional forms and predictions presented here.

In support of this qualitative generality is the fact that the practical totality of the experimental work mentioned above reports decreasing latent periods reaching a minimum plateau as growth rate increases, resembling mildly-declining exponentials, whereas burst sizes increase rapidly. The degree of plasticity for *B* is significantly more marked than that for *L* (e.g. (Webb et al., 1982) reports a 18-fold variation in *B* across different viral strains as *µ* increases). We have observed such behavior under stationarity, with and without associated stochasticity and oscillations, which shows the robustness of these results across environmental conditions.

### The mechanisms underlying the observed latent periods, burst sizes, and trade-offs

The plasticity expressions introduced here shed some light about the mechanisms underlying the plastic behavior of these traits. In our system, the emerging latent period results from the interaction between viral plasticity (encoded in *E*(*µ*)) and the timing for infected-cell removal (the 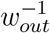 term). The latter shapes *L* across generations and has the potential to reduce the range of variation of *L* with respect to that of *E*(*µ*). When the removal rate is fixed to sufficiently-small values and *µ* varies, the constant term dominates over that of *E*(*µ*), which would lead to the mild effect of plasticity on *L* reported here and in experiments. Higher removal rates, however, would result in a more noticeable plasticity for *L*, especially in the low *µ* limit; low growth rates in this case match high [*C*]_*st*_, thus resonating with past theories about the increasing role of plasticity in *L*_*ESS*_ as host density increases (Wang et al., 1996). For the oscillatory cases, the range of variation for *L* is augmented with respect to *E*(*µ*); unfortunately, our expressions fail to provide an analytical understanding of the underlying mechanisms in this case.

Due to the conspicuous degree of plasticity for the maturation rate, an increase in host performance allows the phage to produce more virions with a shorter lytic cycle, thus breaking the classic trade-off. Therefore, we hypothesize that the importance of plasticity does not reside in the modification of the generation time but rather in the pronounced modification of the offspring number. On the other hand, when controlling the host growth rate by tuning *w*_*out*_, decreasing the dilution rate barely increases *µ* while simultaneously decreases infected-host removal, rendering plasticity negligible: when the survival probability of the offspring is increasingly higher and the abundance of available hosts decreases, it is advantageous to lyse hosts at later times and synthesis time becomes mostly irrelevant. With negligible plasticity, higher burst sizes can only be achieved with longer latent periods, leading to the classic positive correlation between *L*_*ESS*_ and *B*_*ESS*_ and negative correlation between *L*_*ESS*_ and [*C*]_*st*_.

### Plasticity inverts expectations for short-term population dynamics

The wide range of possible burst sizes enabled by plasticity affect the ecological dynamics for both host and phage populations, altering dramatically the expectations built with standard models that neglect plasticity. Differently from non-plastic descriptions, in our model better growth conditions for the host lead to an increased phage population concentration, which increases the pressure upon the host reducing its population, and vice versa. Such population response is somewhat compatible with the “Kill the winner” theory (Winter et al., 2010), as increasing host performance translates into an increased viral regulation. This key qualitative difference between the plastic and non-plastic descriptions remains regardless of the value of *µ* chosen to parametrize the latter (in the cases shown here, using values that would be obtained in a hypothetical standard experiment to measure viral traits, i.e. under host optimal conditions). The reason is that the standard non-plastic model uses a flat line to represent traits that can vary widely with host physiology such as the offspring number. Thus, any static parametrization for the virus necessarily introduces a significant error under dynamic conditions.

### Applicability and limitations

Our expressions obviate the use of different parametrizations to capture different locations or environmental changes for the same host-phage system. They also preclude the use of average parametrizations that, in any case, will not be able to account for dynamic changes in host physiology. Available standard models rely exclusively on host number to assess viral performance. Considering viral plasticity, thus, can alter non-trivially the predictions of these standard models through the dynamic interaction between phage and host performance. For biogeochemical models, such interaction determines key aspects such as primary production. Our expressions for the most dominant phenotype can help improve the predictability of such models, as the two-stage chemostat and inflow and outflow of hosts and nutrients roughly represent a volume of oceanic water. Such flows can also represent those in the human gut, and therefore our expression can help the study of phages infecting the gastrointestinal tract. Like for biofilm phages, space will play in these cases a key role imposing marked resource gradients that emphasize the need to consider plasticity instead of artificial space-specific static parametrization. The relevance of space, nonetheless, also increases the relevance of *k* as the time between adsorption events influences the phage strategy. On the other hand, representing the astonishing viral diversity observed in nature is one of the main challenges to including viruses in large-scale models. Plasticity helps include a dynamic phenotypic diversity for the phage community without the need to invoke any genetic change. Accounting for other realistic details such as host predation mortality (which affects cell removal rates), alternation between lytic and lysogenic infection modes, or a more accurate description of the host requires changing the standard model used here and, therefore, may alter our analytical predictions. Future work will include host plasticity and co-evolution, in which aspects such as cell size and number of receptors (both affecting the adsorption rate) play an important role. From a broader perspective, our work can also advance knowledge on how prey physiology (e.g. nutritional level) can affect the ecological and evolutionary strategy of the predator (e.g. prey selection and/or predatory rates).

As a final remark, a note of caution. Although the qualitative agreement between our *L* and *B* and experimental observations shows that our theory includes the key underlying mechanisms, experiments measuring plasticity in *L* and *B* do not reach stationarity, typically lyse cells artificially and do not necessarily target evolutionary stationary states. Stationarity is not reached in the oceans or other realistic scenarios, cases for which our numerical framework is more suitable. Ultimately, additional experimental information is required in order to improve and generalize our theory across systems, thus contributing to advancing our understanding of host-phage interactions.

## Acknowledgments

The authors would like to thank C. Caceres, M. Heath, P. Suthers, J. Weitz, J. Yin, and A. Zaritsky for helpful discussions; E.W. Birch and P. Golec for providing published data for calculations; and IC1 (University of Granada) for the use of its computational infrastructure. M.C. and J.A.B. were supported by the Marine Alliance for Science and Technology for Scotland (MASTS) pooling initiative, funded by the Scottish Funding Council (HR09011) and contributing institutions.

## Online Appendix

### A Data compilation

In order to deduce expressions for the eclipse period, *E*, and maturation rate, *M*, of the virus as a function of the host growth rate, *µ*, we compiled the available information from (Birch et al., 2012; Golec et al., 2014; Hadas et al., 1997; You et al., 2002). These studies followed one infection cycle of T-phages, with the bacterium *Escherichia coli* as a host. Specifically, the aim of these “one-step growth” experiments was to measure the viral traits for various host growth rates at the moment of infection.

For (Birch et al., 2012; Golec et al., 2014; You et al., 2002), the experimental plaque-forming unit (PFU) data obtained artificially lysing the cells with chloroform, which ultimately can be used to monitor the number of intracellular and extracellular virions, were either provided in the publication or by the authors in personal communications. In such cases, we extracted our own measurements for *E* and *M* for each host growth rate, which allowed us to work with trait values consistently obtained using the same methodology for data extraction. To this end, we used the standard definitions of eclipse period and maturation rate. The former is defined as the time after infection at which the first virion is assembled, whereas the latter can be defined as the slope of the increasing part of the chloroform-generated PFU data, approximated as a linear function of time (You et al., 2002). The resulting points for each growth rate are shown in Figs.A1–A3. Our calculation of such *E* and *M* points agrees, within error bars, with the data points shown in (You et al., 2002) (only publication that provides both experimental data and resulting *E* and *M*). The slight displacement of the whole dataset values observed in Fig.A1 does not affect qualitatively the analysis presented below.

In the case of (Hadas et al., 1997), *E* and *M* values were provided in the publication. A more recent version with an improved methodology to obtain *E* and *M* was presented in (Rabinovitch et al., 1999). The original dataset presented a high level of noise (A. Zaritsky, personal communication), which the newer methodology aimed to ameliorate using moving averages. With that in mind, we used this latter compilation for our analysis, although the use of the older dataset does not change qualitatively the results. Because (Golec et al., 2014) focused on low-growth rates only, we analyzed the data both separately and pooled together with (Hadas et al., 1997; Rabinovitch et al., 1999), which used the same viral strain at the same temperature. Such an exercise allowed us to test our choice of functional forms with increased statistical power, thus enabling a more complete picture of plasticity for the T4 phage.

### B Data analysis

In order to obtain a functional form capturing the relationship between *E* and *M* and host growth rate, *µ*, we explored biologically-reasonable curves able to explain the data above.

From the qualitative point of view, the eclipse period in all cases shows a negative correlation with host growth rate, with a lower plateau for high *µ* (Figs.A1–A3, left panels). The data from (Rabinovitch et al., 1999) and (Golec et al., 2014) seem to connect reasonably well, which suggests a negligible effect of the method to control *µ* upon these one-step growth experiments. Decreasing exponentials match such qualitative behavior across databases and are conceptually simple. A possibility with a reduced number of parameters is:

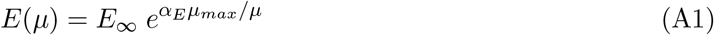

where, to facilitate comparison across examples, we normalized *µ* using the maximum growth rate (*µ*_*max*_) reported in the nutrient-rich or highest dilution rate case, in all cases compatible with tabulated values for *E. coli* maximum growth rate at the temperature used in each experiment (Herendeen et al., 1979). Such normalization only redefines the *α*_*E*_ parameter and therefore does not influence qualitatively our results. Eq.(A1) assumes that *µ* = 0 for the host leads to an infinite *E*, effectively preventing viral reproduction. However, in reality reproduction is still possible for very low growth rates (Golec et al., 2014), whereas extremely-low growth rates may lead the virus to choose a different host or enter a lysogenic mode. In addition, such function does not capture the apparent plateau that occurs for high growth rates. A more suitable option is:

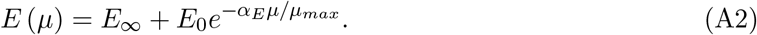

Although this function adds a parameter, it is conceptually easier to understand, as there is a clear minimum-value plateau and the function can capture the higher (but not infinite) *E* values observed for very low growth rates. In order to test the closeness of the proposed functions to the data, we fitted each database using the Matlab Curve Fitting Toolbox with the Trust-region algorithm and all the default options, using our custom functions for the fitting. Eq.(A2) explained the eclipse period data compiled for all databases, including the pooled data for the T4 phage (table A1). Although somehow considered in statistics such as the adjusted *R*^2^, the fact that there are few points and three parameters contributes to the unusually-high value for the goodness of fit indicators (the adjusted *R*^2^ and the root-mean-squared deviation) at the expense of particularly-high errorbars for some of the obtained parameters. In some cases, we forced the fitted curve to pass through the *E*(*µ* = *µ*_*max*_) value of the database, effectively defining *E*_∞_ and thus reducing the number of parameters. All such possibilities just change slightly the shape of the curve, that retains the main features observed in the experiments (decreasing trend with apparent plateau), and do not affect our ecological and evolutionary analyses.

On the other hand, the maturation rate curve shows an initial slow growth that accelerates for intermediate growth rates, followed by an apparent saturation for at least two of the three data compilations for which high *µ* values are available (Figs.A1–A3, right panels). This qualitative behavior resembles that of a sigmoid function, for example:

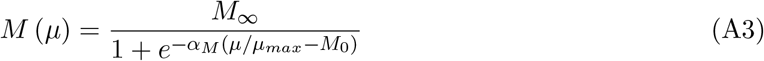

Both chosen forms for *E*(*µ*) and *M* (*µ*) have a biological interpretation rooted in how changes in host growth affect its synthesis machinery and rates (see main text), interpretation that is supported by how temperature and T-phage type affects the *E* and *M* curves (see Fig.A4). For *E*, increasing temperature reduces the differences between synthesis times for low and high growth rate and consequently decreases the minimum time plateau (suggesting an increase in the overall efficiency of synthesis). Moving from T7 to T4 reduces significantly such performance as T7 combines early on host RNAP with its own (much more efficient) for the synthesis of proteins (Molineux, 2006). The closeness of the T7 at 30*C* curve and the T4 curve may indicate a compensating effect of temperature over host RNAP performance for T4. In the case of *M*, increasing temperature leads to the consequent overall increase in the assembly rate, whereas moving from T7 to T4 increases significantly the saturation value; with caution due to the quality of the T4 dataset (see above), the latter may be interpreted as a bottleneck relief resulting from the dependence of both parts of T4 virion production (synthesis and assemblage) on the host machinery.

Other functions may also fit existing data. The only set of functional forms describing the link of *E* and *M* with host quality currently available in the literature (Rabinovitch et al., 2002) is case-specific and does not hold across examples (i.e. they cannot explain the qualitative behavior observed for other systems). For instance, their proposed functional form for *E* does not show the lower plateau, and is non-monotonic thus leaving unexplained the increase in *E* for low growth rates reported for the same phage under the same temperature in (Golec et al., 2014). Their proposed *M* curve does not show any saturation, which contrasts with the expected saturating behavior observed for metabolic rates as growth conditions improve (You et al., 2002). Nonetheless, as explained below our results are robust against the specifics of the chosen functional forms for *E*(*µ*) and *M* (*µ*).

### C Model analysis: ecological long-term behavior

The ecological dynamics of the model are described by Eqs.(3)–(6), with *L* being defined as the main adaptive trait of the virus and *B* provided by the trade-off function, Eq.(7). The environmental conditions are set up using a two-stage chemostat, which enables ecological stationary states in the long term by providing an inflow of fresh hosts and nutrient, and an outflow that removes free and infected hosts, free viruses, and nutrients. The addition of fresh cells also reduces the pressure on the host, which in turn reduces the occurrence of the “predator-prey-like” population-level oscillations typically observed in host-phage models, and the probability for the host to engage in a co-evolutionary arms race with the phage. The latter, thus, facilitates an evolutionary steady state.

For simplicity, we assume the adsorption rate to be constant, as here we do not consider many of the factors that can affect this trait (see main text). Other assumptions are *i)* both chemostats are well stirred, and therefore in the second chemostat host and phage encounter each other randomly; *ii)* multiple infections do not occur; *iii)* the host stops nutrient uptake at the moment of infection, and the synthesis machinery only focuses on viral replication.

From the ecological point of view, in the long term the system will reach a stationary state, which is mathematically equivalent to imposing *d*[*C*]/*dt* = *d*[*I*]/*dt* = *d*[*V*]/*dt* = *d*[*N*]/*dt* = 0. Such a condition leads to *i)* a trivial state in which [*V*]_*st*_ = [*I*]_*st*_ = 0, [*C*]_*st*_ = *w*_*in*_[*C*_0_]/(*w*_*out*_ − *µ*_*st*_), or *ii)* the non-trivial state where virus and host co-exist:

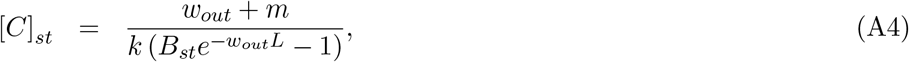

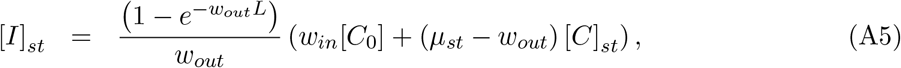

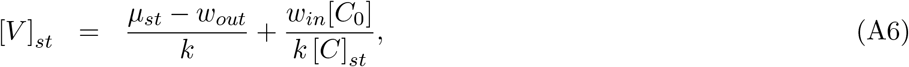

where the nutrient concentration has also reached its stationary value, [*N*]_*st*_. Such a value determines the stationary host growth rate, *µ*_*st*_ = *µ*_*max*_[*N*]_*st*_/([*N*]_*st*_ + *K*_*N*_) and, therefore, *B*_*st*_ through *E* and *M* (Eq.(7)). The dependence of *E* and *M* on *µ*([*N*]), however, prevents a closed analytical expression for [*N*]_*st*_. This non-trivial stationary state is feasible as long as *L* < ln(*B*_*st*_)/*w*_*out*_ and *µ* > *w*_*out*_ − *w*_*in*_[*C*_0_]/[*C*]_*st*_. On the other hand, we point the reader to (Bonachela and Levin, 2014) and references therein for the discussion on the stability of this solution. Note that the minimum possible growth rate for which the non-trivial stationary state is stable is given by *µ*_*st*_ = *w*_*out*_; below such growth rate, the system falls into the trivial stationary state.

### D Model analysis: first chemostat

As pointed out above, the first chemostat aims at providing a constant flow of host cells to the second chemostat. Thus, [*C*_0_] corresponds to the stationary concentration of hosts in the first chemostat. The stationary nutrient concentration in such a chemostat is given by [*N*_1*st*_] which, together with an additional amount [*N*_*ext*_] provided by an external source, compose the total input nutrient for the second chemostat, i.e. [*N*_0_] = [*N*_1*st*_]+[*N*_*ext*_]. The external source of nutrient, [*N*_*ext*_] aims at enabling control over [*N*_0_] without altering the first chemostat. For simplicity, we assume that [*N*_*ext*_] goes into the second chemostat at a rate *w*_*in*_. On the other hand, because the outflow rate of the first chemostat represents the inflow rate of the second chemostat, the stationary state of the former is that of a regular chemostat with a dilution rate *w*_*in*_, provided by:

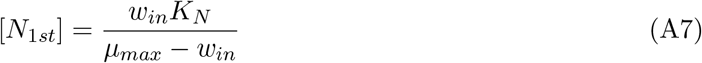

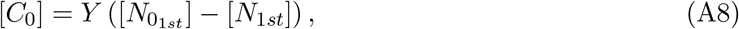

where 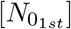 is the source of nutrients for this first chemostat. Tuning *w*_*in*_ determines, therefore, the inflow of nutrients from the first chemostat into the second. Furthermore, we can fine-tune [*C*_0_] by altering 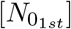, thus allowing for the control the concentration of fresh hosts independently from [*N*_1*st*_]. The limit [*C*_0_] = [*N*_1*st*_] = 0 transforms the system in a classic one-stage chemostat.

### E Model analysis: evolutionary long-term behavior

To reach an analytical expression for the evolutionary steady state, we conducted invasion analyses. An invasion analysis provides the strategy that maximizes the fitness for an evolving agent within a dynamical system, i.e. the evolutionarily stable strategy (ESS). In our case, phages are the only evolving agent. From the viral traits defined in the main text, *k* is assumed to be constant (see above), and *E* and *M* are set by the host growth rate through plasticity, Eqs.(A2) and (A3). *B* is determined by *M*, *E*, and *L* through a trade-off, traditionally assumed to be linear (Eq.(7)). Thus, *L* is the only remaining trait and, therefore, it characterizes the phenotype, which in turn identifies each phage population. Because the holin gene controls the latent period (see main text), finding the viral genotype that maximizes fitness translates into finding the most competitive *L*.

In an invasion analysis, the stability of a system in which an invader/mutant attempts to out-compete a resident phage is studied. It is assumed that *i)* the phenotype winning such competition completely excludes the other, and *ii)* every mutant enters the system in a small amount and only when the resident viral phenotype and host have reached their ecological stationary state. An implicit assumption is that the feasibility and stability conditions of the non-trivial ecological state are fulfilled by the resident.

The invasion analysis is, thus, basically a perturbation analysis in which we study the stability of a system in which a mutant phenotype, characterized by *L* = *L*_*M*_, challenges the equilibrium for the viral resident and host. If the system is unstable, the mutant outcompetes the resident, and vice versa. Sampling any combination of mutant and resident, the ESS is the *L*_*R*_ for which any competing mutant shows a non-positive fitness, i.e. the system is stable (alternatively, any mutant *L*_*M*_ whose fitness is never negative when competing against any possible resident, i.e. the system is unstable).

Thus, once the resident and the host have reached stationarity, we study the stability of the system of equations:

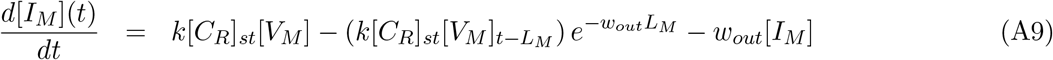

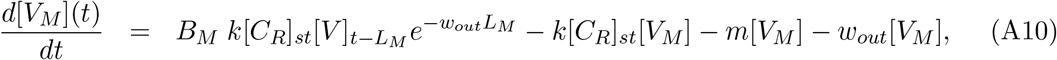

where [*C*_*R*_]_*st*_ is obtained from Eq.(A4) with *L* = *L*_*R*_. Note that, because [*C*_*R*_]_*st*_ is constant, the host growth rate does not play any role in the invasion analysis.

To study the stability of the system, we need to study the eigenvalues *λ* associated with Eqs.(A9)–(A10), for which we need to solve the characteristic equation 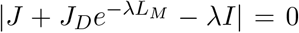 (Beretta and Kuang, 2001). *J* represents the Jacobian matrix associated with the instantaneous terms of the equations, *J*_*D*_ that of the delayed terms, and *I* the identity matrix. Such an equation, in our case, is given by:

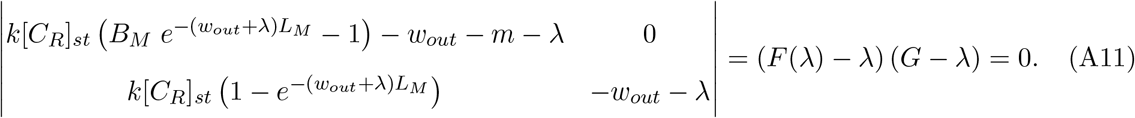

A trivial solution is given by *λ*_1_ = *G* = −*w*_*out*_, and the other solution given by 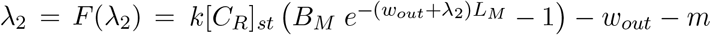. Because *λ*_1_ is negative, the stability of the system is determined by the sign of *λ*_2_. Thus, we focus on the condition *λ*_2_ = *F* (*λ*_2_) = 0, that is:

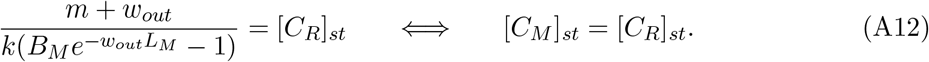

If [*C*_*R*_]_*st*_ < [*C*_*M*_]_*st*_, *λ*_2_ < 0 and the resident resists invasion (alternatively, the mutant invades if [*C*_*M*_]_*st*_ < [*C*_*R*_]_*st*_). Therefore, like in classic competition theory Tilman (1982), the ESS is given by the strategy minimizing the host concentration (only viral resource), i.e. the strategy that is solution to the equation *d*[*C*]_*st*_/*dL* = 0, which ultimately translates into the equation *dB/dL* = *w*_*out*_ *B* (see Eq.(A4)). Using the trade-off equation for *B*, Eq.(7), we finally reach the expression for the ESS, given by:

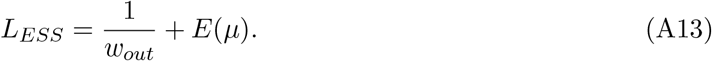

See (Bonachela and Levin, 2014) for further details, including other forms for the trade-off, and references. Note that the plastic potential of the virus is enclosed in the eclipse period, *E*. Using Eq.(A13) together with Eq.(7) provides an expression for burst size associated with this ESS:

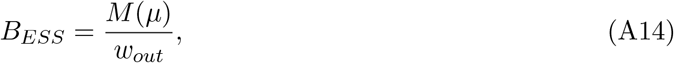

which also shows plasticity potential, now through the maturation rate. Due to the assumption of quick ecological stationarity for the host during the deduction, this expression and that of the non-plastic case (*L*_*non*_ and *B*_*non*_, respectively) are very similar. Indeed, and following the discussion in the main text, we can obtain the expression for *L*_*non*_ and *B*_*non*_ by using the maximum growth rate measured in the corresponding laboratory or *in silico* experiment together with Eq.(A13) and (A14), respectively. For cases in which the removal rate for infected cells (here, dilution rate, *w*_*out*_) is constant, the ESS of the non-plastic case coincides with the minimum value for the plastic *L*_*ESS*_ and maximum for *B*_*ESS*_ (see Fig.2). When *w*_*out*_ is fine tuned to alter the host growth rate, however, *L*_*non*_ and *B*_*non*_ become curves when represented against *µ* (Fig.3), as the environmental fine-tuning affects the first term of the *L*_*non*_ and *B*_*non*_ expressions above. The differences with *L*_*ESS*_ and *B*_*ESS*_ are very small in this case because plasticity becomes relevant only in the limit of very high dilution rates (high mortality for the population).

We can further characterize the ESS of the plastic virus by performing a numerical invasion analysis for every possible pair resident-mutant. Plotting the sign of the fitness of the mutant (or invasion eigenvalue, *λ*_2_) enables the graphical determination of the ESS, or resident strategy for which any mutant shows non-positive fitness. Such representation, the pairwise invasibility plot (PIP), also provides information about the evolutionary convergence of the ESS Geritz et al. (1998). In Fig.A5 (left panel), for example, we can observe the agreement between the ESS obtained from Eq.(A13) and that obtained graphically. Moreover, such point is convergence-stable, as phenotypes closer to the ESS can always invade populations formed by phenotypes further from *L*_*ESS*_. The host growth rate was calculated numerically for each resident latent period by integrating the equations to stationarity.

### F Model simulations: eco-evolutionary long-term behavior

We can also determine the ESS numerically by using an eco-evolutionary framework in which the constraints and assumptions of the invasion analysis are relaxed. Such a framework has been successfully used in the past to study the long-term behavior of systems where both phenotypic plasticity and evolution occurred simultaneously and therefore had the potential to interact (e.g. (Bonachela and Levin, 2014; Lomas et al., 2014)).

Starting from an initial clonal phage population and a clonal host population, Eqs.(3)–(6) are integrated numerically thus providing the ecological dynamics of the host-phage system. At exponentially-distributed random times (or, indistinctly, periodically), new mutants are introduced in the system. The frequency of such an event sets the effective mutation/invasion rate, with the low frequency (or large period) limit approximating the invasion analysis situation above. Mutants result from a genetic algorithm in which mutant phage populations are created from randomly-chosen parents; the mutant’s latent period results from a random variation within a 20% of that of the parental population. The probability for a phenotype to be chosen for mutation is provided through a roulette-wheel selection process by which the higher the population density, the higher the probability to be selected. New mutants enter the system in small amounts forming a new clonal population that is described by Eqs.(A10)–(A9), equations that are added to the existing system of equations for numerical integration. Iterating the process, ecological interactions and evolution occur simultaneously, with mutants entering the system and others going extinct via natural selection as they all compete for the same single resource (the host). In this way, there is an alternation of the dominant phenotype that can be visualized by plotting the dominant *L* in the system against time (Fig.A5, right panel). Eventually, a phenotype arises that cannot be challenged by any other phenotype under the particular conditions of the system and, therefore, such a phenotype remains dominant until the final time imposed in the simulation. Such a phenotype is the stationary state in the dominant-*L* versus time representation, but also in the representation of the average *L* of the total phage population versus time, as the phenotype with *L*_*ESS*_ overpowers such average. Because mutation (represented with the genetic algorithm) is an intrinsically-stochastic way to explore the phenotypic space, different replicates may reach evolutionary stationary states that are close but not necessarily exactly at the ESS (see Fig.A5, right). The ESS can be found as the average over replicates of the stationary *L* values.

In addition to the results discussed here and in the main text, this framework has allowed us to go beyond the limitations of experimental observations and show that, for example, there is a limit as to how much the burst size can grow with host growth rate, or that the maturation period (period elapsing from eclipse to the end of the latent period, mathematically *L* − *E*, in our model 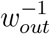) does not depend on the host growth rate.

Our analytical predictions matched those of the numerical framework for all the evolutionary analyses in which a stationary state could be defined (see main text). Such cases also include stochastic stationary states in which the ecological interactions fluctuate randomly around a well-defined and most-probable value on average. Indeed, similar results were obtained when allowing [*N*_*ext*_] only or both [*N*_*ext*_] and [*N*_1*st*_] to vary randomly by a factor *σ* = 0.3 or *σ* = 0.5 (Fig.A6, right). The analytical expressions failed to describe the emerging ESS only for the cases for which the one-stage chemostat showed wide oscillations that translated into ecological oscillations for, e.g. the host population that form a very skewed distribution (*µ/µ*_*max*_ > 0.35 in Fig.A6, left). Regardless of the agreement with Eq.(A13), the emerging *L*_*ESS*_ decreased with the host growth rate and *B*_*ESS*_ increased when using [*N*_0_] or *K*_*n*_ to control *µ*; both *L*_*ESS*_ and *B*_*ESS*_ increased when using the dilution rate to increase *µ*.

### G Model simulations: remineralization

When exploring the predictions of the model for the three consecutive dynamic events, we tested whether remineralization of the cellular content of the lysed cells would change in any way the observed pattern. To this end, we implemented a version of the model with a simplistic +*R Q* (*k*[*C*]_*t*−*L*_[*V*]_*t*−*L*_) *e*^−*w*^ term in Eq.(6), with *R* the remineralization rate and *Q* the nutrient content per cell. We did no observe any qualitative change with respect to the plastic case (not shown) and, therefore, we conclude that considering remineralization does not alter any of the trends reported here.

### H Additional Results

[Figures A7–A9 here]

---

**Table A1:** Parameters obtained with the chosen functional forms for *E* and *M*, Eq.(1) and (2).

|  | You | Birch | Hadas+Golec |
| --- | --- | --- | --- |
| $E_\infty$ ($min$) | 15.33(2.38) | 9.16(1.42) | 18.26(3.58) |
| $E_0$ ($min$) | 82.71(34.29) | 14.00(12.6) | 94.18(25.12) |
| $\alpha_E$ | 5.9(1.0) | 6.62(4.30) | 7.76(4.15) |
| Adj. $R^2$ for $E$ fit | 0.99 | 0.98 | 0.89 |
| NRMSD | 0.017 | 0.063 | 0.109 |
| $M_\infty$ ($vir. min^{-1}$) | 6.7(0.7) | 23.14(13.75) | 47.32(6.82) |
| $M_0$ | 0.64(0.09) | 0.68(0.31) | 0.65(0.04) |
| $\alpha_M$ | 9.70(8.45) | 4.33(2.49) | 12.03(2.83) |
| Adj. $R^2$ for $M$ fit | 0.98 | 0.99 | 0.99 |
| NRMSD | 0.076 | 0.017 | 0.027 |

The number between parentheses indicates the error associated with the parameter (one standard deviation). Note that we normalized the root mean-squared deviation using the maximum range of variation of the dependent variable, thus obtaining a fractional value that can be compared across fits (NRMSD).

**Table A2:** Symbols for variables used in the model and parameter values.

| Symbol | Description | Units | Value |
| --- | --- | --- | --- |
| $[N]$ | Dissolved Inorganic Nitrogen Concentration | $mol\ l^{-1}$ | Ecological variable |
| $[C]$ | Non-infected Host Concentration | $cell\ l^{-1}$ | Ecological variable |
| $[I]$ | Infected Host Concentration | $cell\ l^{-1}$ | Ecological variable |
| $[V]$ | Free Virus Concentration | $cell\ l^{-1}$ | Ecological variable |
| $M$ | Maturation Rate | $virions\ d^{-1}$ | Eq.(2) and table A1 |
| $E$ | Eclipse Period | $d$ | Eq.(1) and table A1 |
| $L$ | Latent Period | $d$ | Evolutionary variable |
| $B$ | Burst Size | $virions$ | Evolutionary variable |
| $\mu_v$ | Virus Population Growth Rate | $d^{-1}$ | Variable |
| $\mu$ | Host Population Growth Rate | $d^{-1}$ | Variable |
| $\mu_{max}$ | Maximum Host Population Growth Rate | $d^{-1}$ | 40.8 |
| $Y$ | Yield Parameter | $cell\ mol^{-1}$ | $9 \times 10^{13}$ |
| $K_N$ | Half-Saturation Constant the Nutrient | $mol\ l^{-1}$ | $10^{-6} - 2 \times 10^{-4}$ |
| $k$ | Adsorption Rate | $l\ cell^{-1}\ d^{-1}$ | $9 \times 10^{-10}$ |
| $m$ | Virus Mortality Rate | $d^{-1}$ | 0.09 |
| $[C_0]$ | Non-infected Host Supply Concentration | $cell\ l^{-1}$ | 0; $10^8$ |
| $[N_0]$ | Dissolved Inorganic Nutrient Supply Concentration | $mol\ l^{-1}$ | $45 - 2000 \times 10^{-6}$ |
| $w_{out}$ | Chemostat Dilution Rate | $d^{-1}$ | 0.24 – 21.84 |
| $[N_{ext}]$ | External Source for Dissolved Inorganic Nutrient | $mol\ l^{-1}$ | From $[N_0] = [N_{1st}] + [N_{ext}]$ |
| $[N_{1st}]$ | Dissolved Inorganic Nutrient in First Chemostat | $mol\ l^{-1}$ | Eq.(A7) |
| $[N_{01st}]$ | Dissolved Inorganic Nutrient Supply First Chemostat | $mol\ l^{-1}$ | From Eq.(A8) |
| $w_{in}$ | Chemostat Inflow Rate | $d^{-1}$ | $w_{out}$ |

Data for the host obtained from (Fagerbakke et al., 1996; Herendeen et al., 1979; Schulze and Lipe, 1964); data for the virus into the ranges shown/used in (Bull et al., 2006; De Paepe and Taddei, 2006; Shao and Wang, 2008; Weitz et al., 2005). For the calculation of the yield factor from the references, we assumed a fixed carbon content per host
cell of $10^{-12}\ g$ .

## Online appendix figures

**Figure A1:**
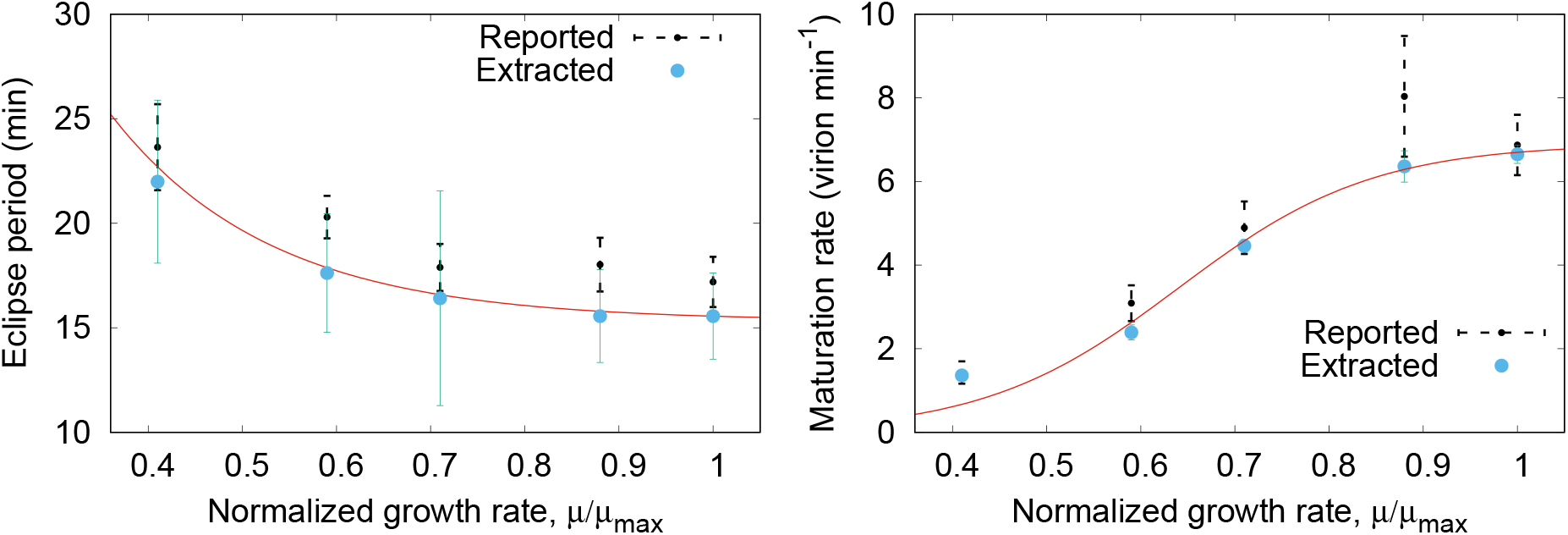
Data extracted from the original one-step growth experiment data in (You et al., 2002) (big dots), values provided in the publication (dashed lines), and our proposed curve (solid line, see fit results in table A1). Left: Eclipse period, *E*; Right: Maturation rate, *M*.

**Figure A2:**
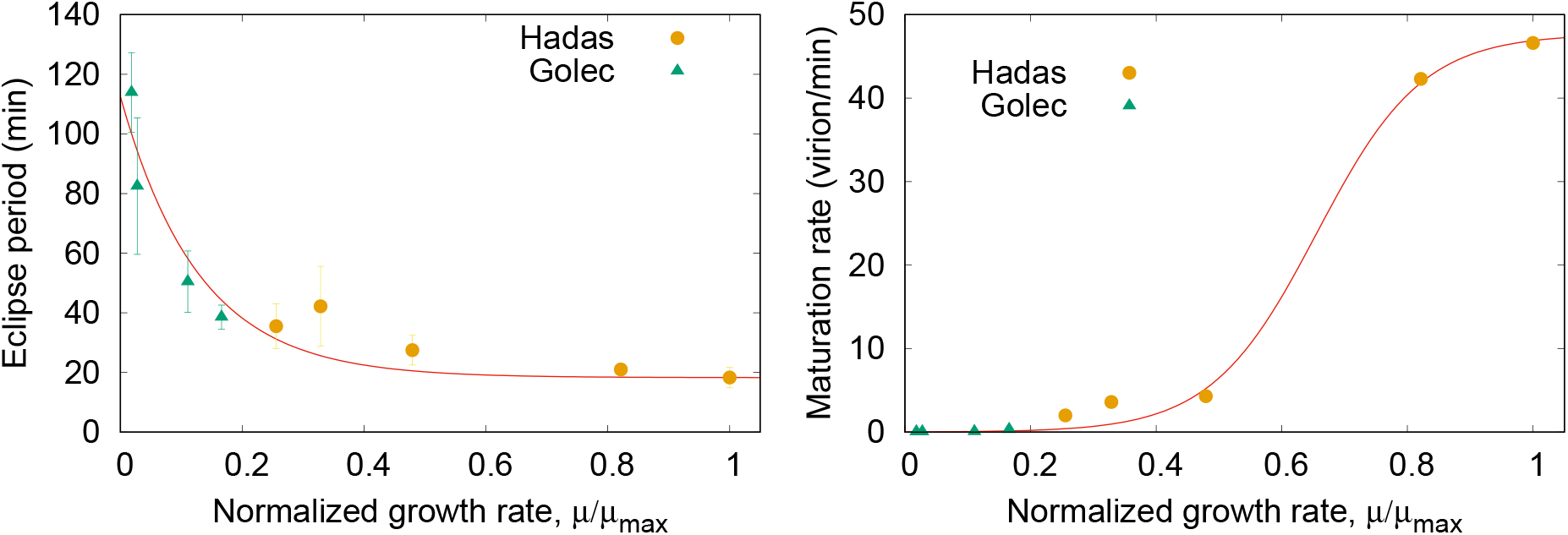
Data from the original one-step growth experiment data in (Golec et al., 2014) and reported data from (Rabinovitch et al., 1999); our proposed curve results from fitting the two datasets together (table A1). Left: Eclipse period, *E*; Right: Maturation rate, *M*.

**Figure A3:**
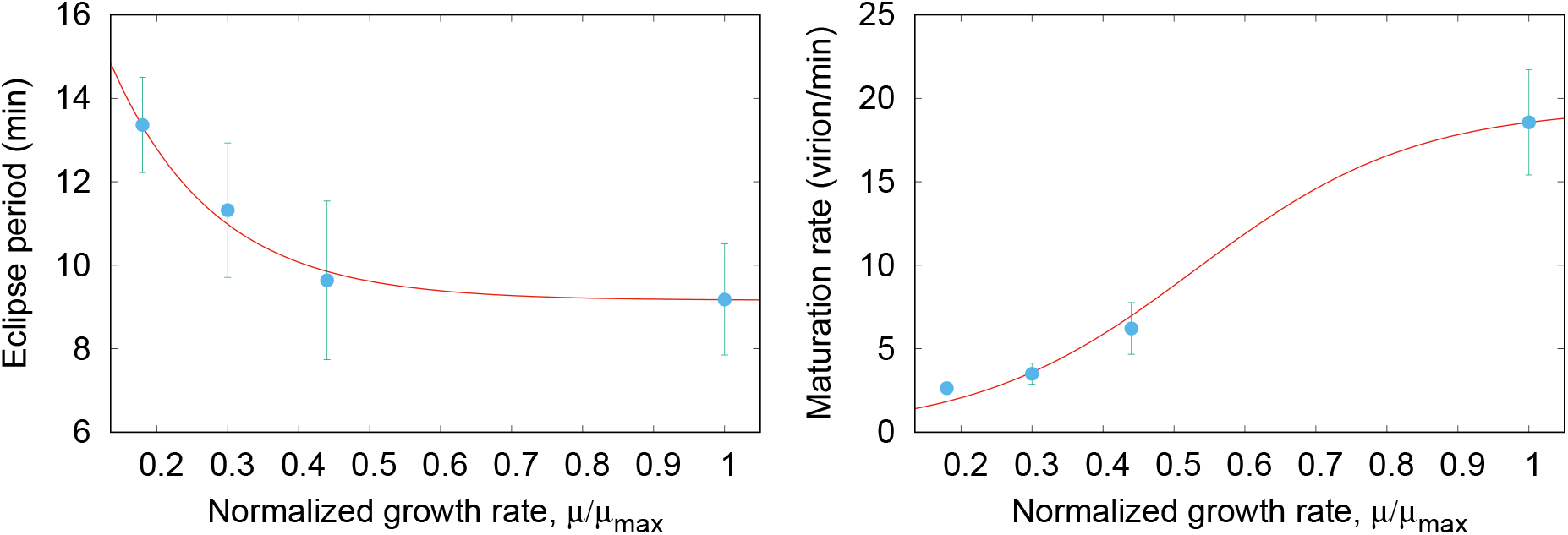
Data extracted from the original one-step growth experiment data in (Birch et al., 2012); the curve corresponds to our fit (table A1). Left: Eclipse period, *E*; Right: Maturation rate, *M*.

**Figure A4:**
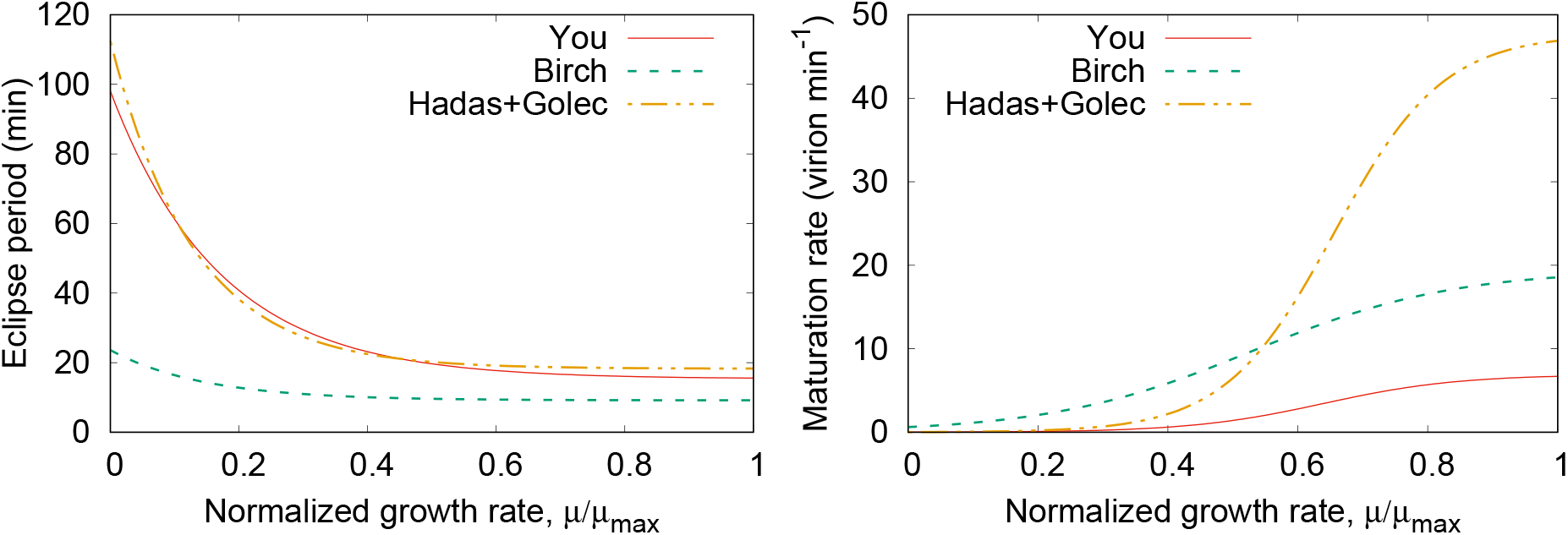
Compilation of the three data-informed curves obtained for *E* (left) and *M* (right) which, together with tables 1 and A1, helps understand the role of temperature and viral species on viral plasticity.

**Figure A5:**
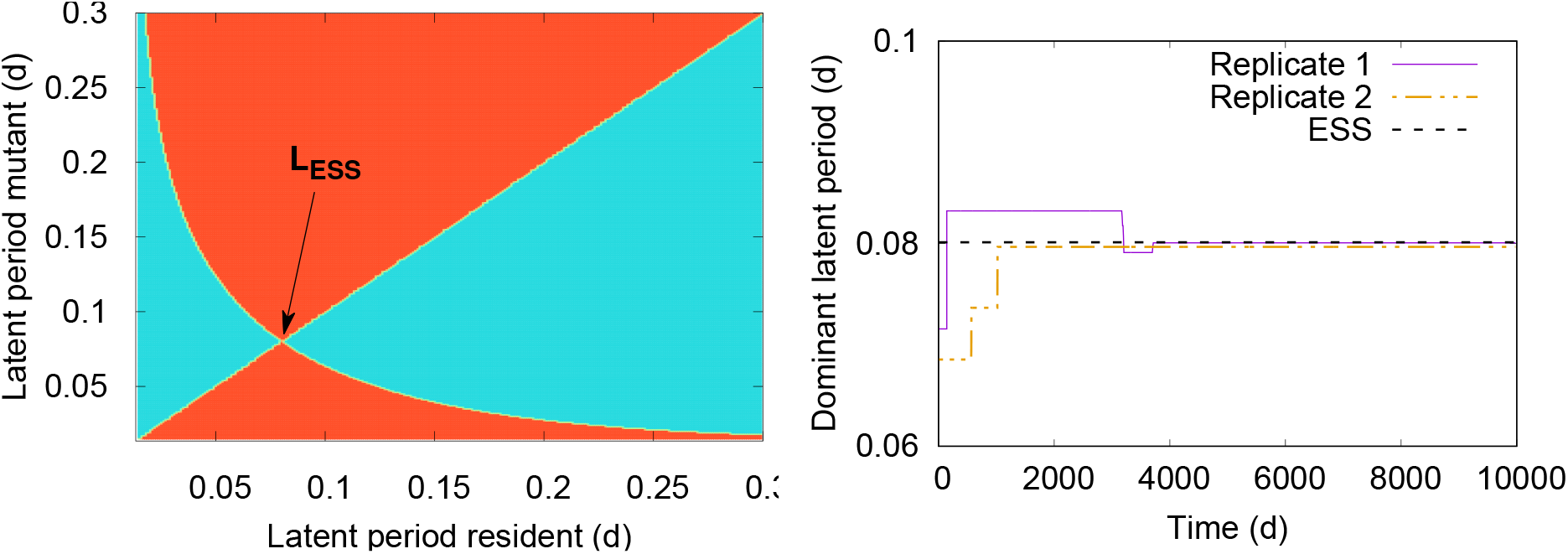
Evolutionary results for the model when Ε(μ) and *M*(μ) are parametrized to fit the data in (You et al., 2002). Left: PIP associated with *w*_*out*_ = 15 *d*^−1^ and *μ* = 20.84 *d*^−1^, for which Eq.(8) predicts *Less* = 0.0801 d; red areas represent negative mutant fitness whereas blue areas represent positive fitness. Right: Evolutionary path in two different replicates for [*N*_0_] = 10^−4^*mol l*^−1^ and *w*_*out*_ = 15*d*^−1^, showing the convergence to the ESS by alternation in the dominant phenotype; *B* shows a similar road to reach *B*_*ESS*_ ~ 146 *virions host*^−1^.

**Figure A6:**
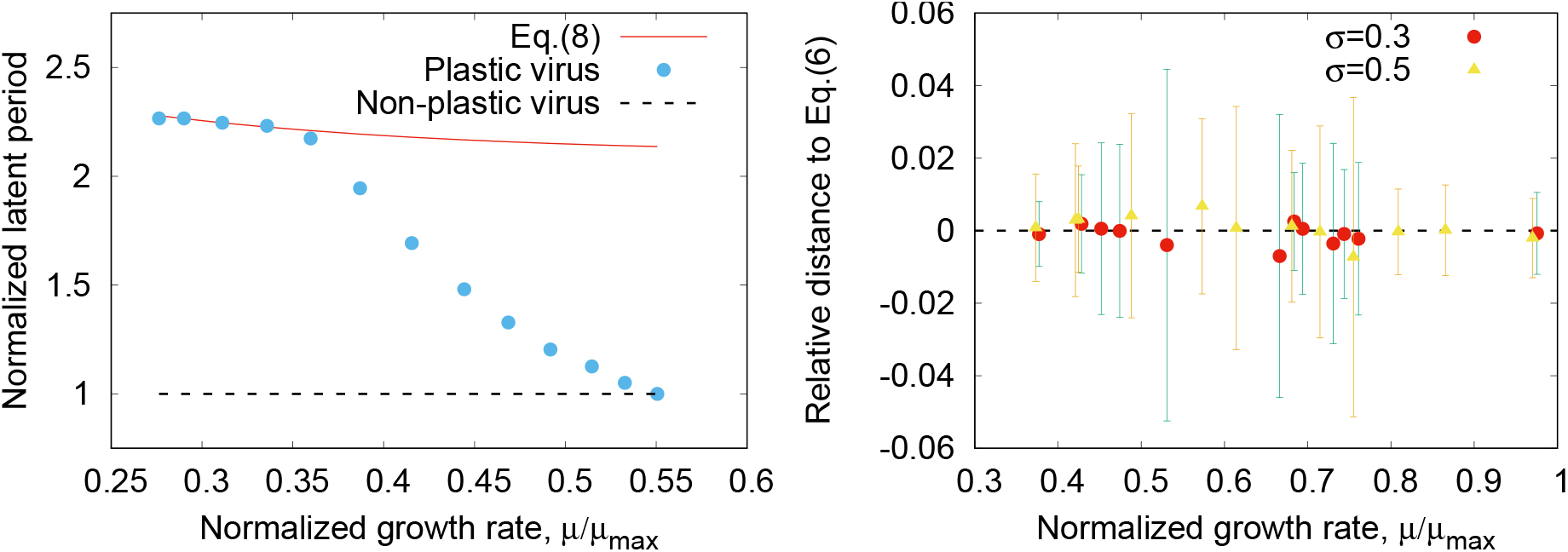
ESS obtained in highly-changeable environments using the input concentration to set the host growth rate. Left: In a one-stage chemostat with *w*_*out*_ = 8.16 *d*−1; the points that depart from predictions match the cases for which oscillations were observed. Right: In a two-stage chemostat with [*N*_*ext*_] changing daily randomly up to a factor *σ* = 0.3 (circles) or *σ* = 0.5 (triangles).

**Figure A7:**
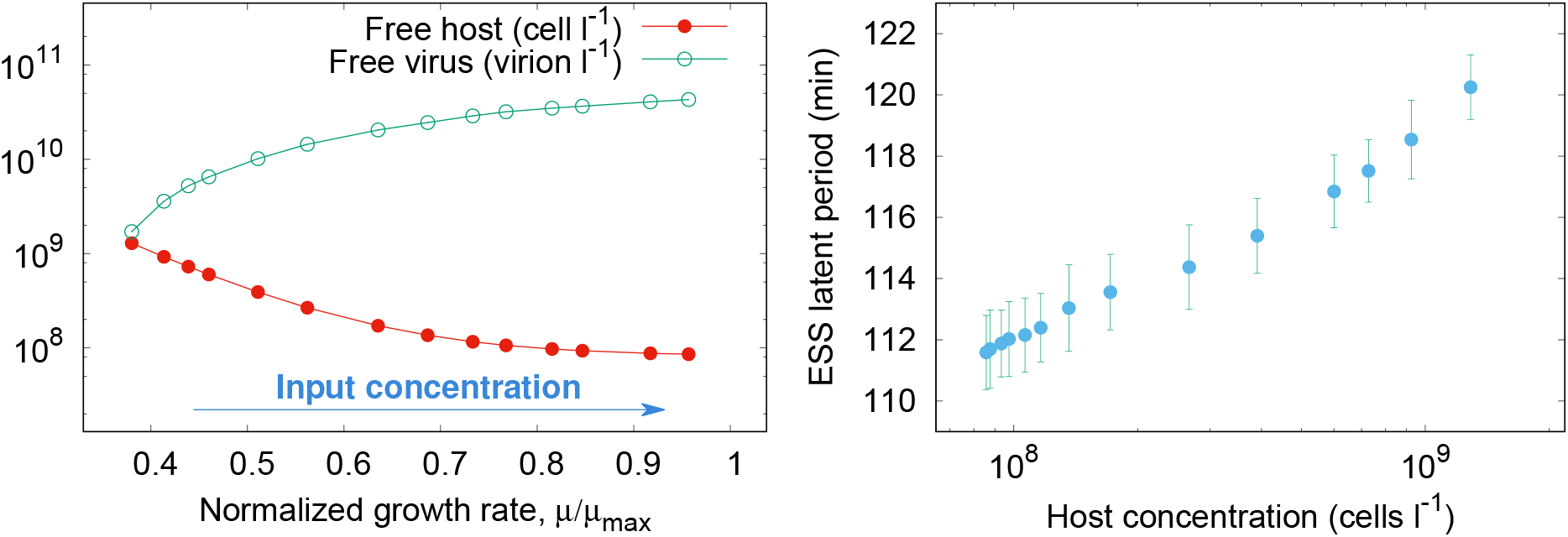
Left: Host and viral concentration as a function of the normalized growth rate (positively correlated with [*N*_0_]) when [*N*_0_] is tuned to control *µ*. Right: Emergent *L*_*ESS*_ as a function of the availability of hosts, [*C*]_*st*_ for the same case.

**Figure A8:**
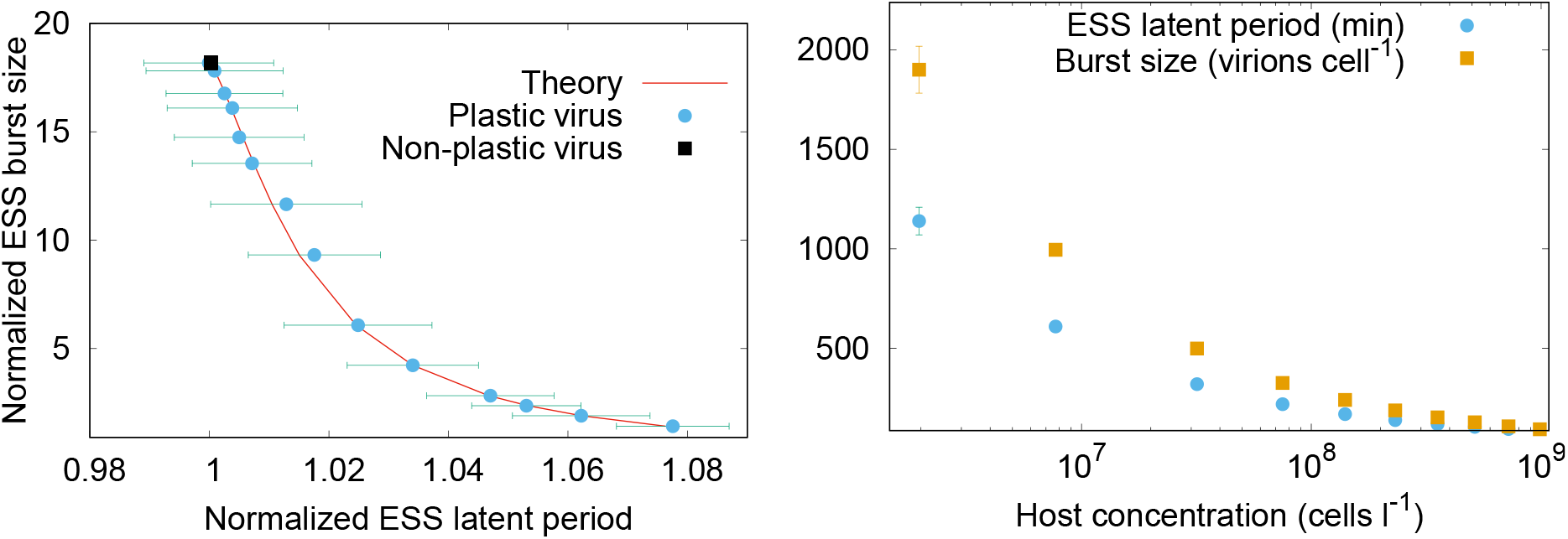
Left: Trade-off curve obtained when host growth rate is increased by increasing [*N*_0_] (see main text for simulation details). Right: Emergent *L*_*ESS*_ and *B*_*ESS*_ as a function of the availability of hosts, [*C*]_*st*_ when the host growth rate is increasing by decreasing the dilution rate.

**Figure A9:**
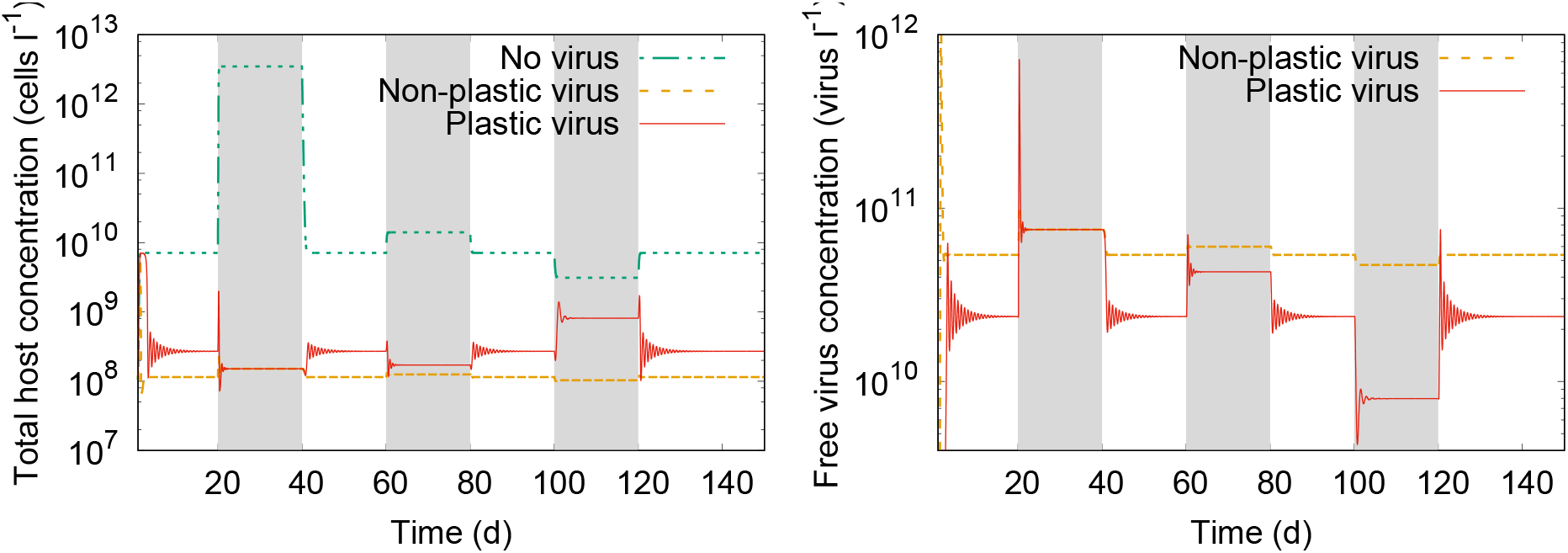
One replicate of the two-stage chemostat in which two host-growth increasing events are followed by nutrient scarcity (shaded areas), as described in the main text. Left: Total number of hosts, including infected (i.e. non-growing) cells. Right: Free phage.

